# Creating De Novo Overlapped Genes

**DOI:** 10.1101/2022.02.15.480625

**Authors:** Dominic Y. Logel, Paul R. Jaschke

**Affiliations:** School of Natural Sciences, Macquarie University, Sydney 2109, New South Wales, Australia

**Author notes:** Correspondence and requests for materials should be addressed to PRJ.

**Keywords:** Deep Learning, Machine Learning, Generative Model, Markov Random Field, Overlapping Genes, Multiple Sequence Alignments, Protein Design, Genome Compression, Synthetic Genomes, Synthetic Biology

## Abstract

Future applications of synthetic biology will rely on deploying engineered cells outside of lab environments for long periods of time. Currently, a significant roadblock to this application is the potential for deactivating mutations in engineered genes. A recently developed method to protect engineered coding sequences from mutation is called Constraining Adaptive Mutations using Engineered Overlapping Sequences (CAMEOS). In this chapter we provide a workflow for utilising CAMEOS to create synthetic overlaps between two genes, one essential (*infA*) and one non-essential (*aroB*), to protect the non-essential gene from mutation and loss of protein function. In this workflow we detail the methods to collect large numbers of related protein sequences, produce multiple sequence alignments (MSAs), use the MSAs to generate Hidden Markov Models and Markov Random Field models, and finally generate a library of overlapping coding sequences through CAMEOS scripts. To assist practitioners with basic coding skills to try out the CAMEOS method, we have created a virtual machine containing all the required packages already installed, that can be downloaded and run locally.

## 1 Introduction

Synthetic biology has led to an explosion in designed genomic parts driving the production of novel functions and molecules [1]. This is done through the construction of genetic circuits with natural or engineered genes controlled by regulatory elements [2]. To make the design of engineered genomes easier, most genome design approaches seek to refactor genomes to remove genetic overlaps and cryptic regulation [3-8], however this does not necessarily provide evolutionary stability to designs[9]. In fact, engineered genes and synthetic architectures often place a deleterious growth burden on the expression host [10-12]. Thus, hosts which have lost the engineered gene have a growth advantage, and over time become the dominant population. This phenomenon has led to ways to constrain evolution by tying the expression of the engineered part to the expression of an essential host component, thus linking organism survival to the retention of the engineered component [13,14]. Design stability is crucial as many future applications for synthetic biology technologies are predicated on usage outside laboratories, such as engineered nitrogen fixation in cereal endophytes [15] and cleaning environmental pollutants [16,17].

A novel way to add genetic stability to engineered genomes is called Constraining Adaptive Mutations using Engineered Overlapping Sequences (CAMEOS) which seeks to emulate the condensed and overlapped coding sequence architecture found primarily in bacteriophage and bacteria [3,18,4,7,19]. This computational approach uses Hidden Markov Models (HMMs) and Random Markov Fields (MRFs) to determine protein residue diversity at a given position, and residue-residue contacts across the proteins, to generate overlapped coding sequences containing two proteins [20].

The foundation for the creation of protein generative models (HMMs and MRFs) is a multiple sequence alignment (MSA) [21]. For the accurate production of these models the MSA must encompass thousands to tens of thousands of sequences (Figure 1A). There are multiple algorithms available for performing protein alignments such as ClustalW [22], FAMSA [23], and MAFFT [24] all of which perform differently.

**Figure 1.**
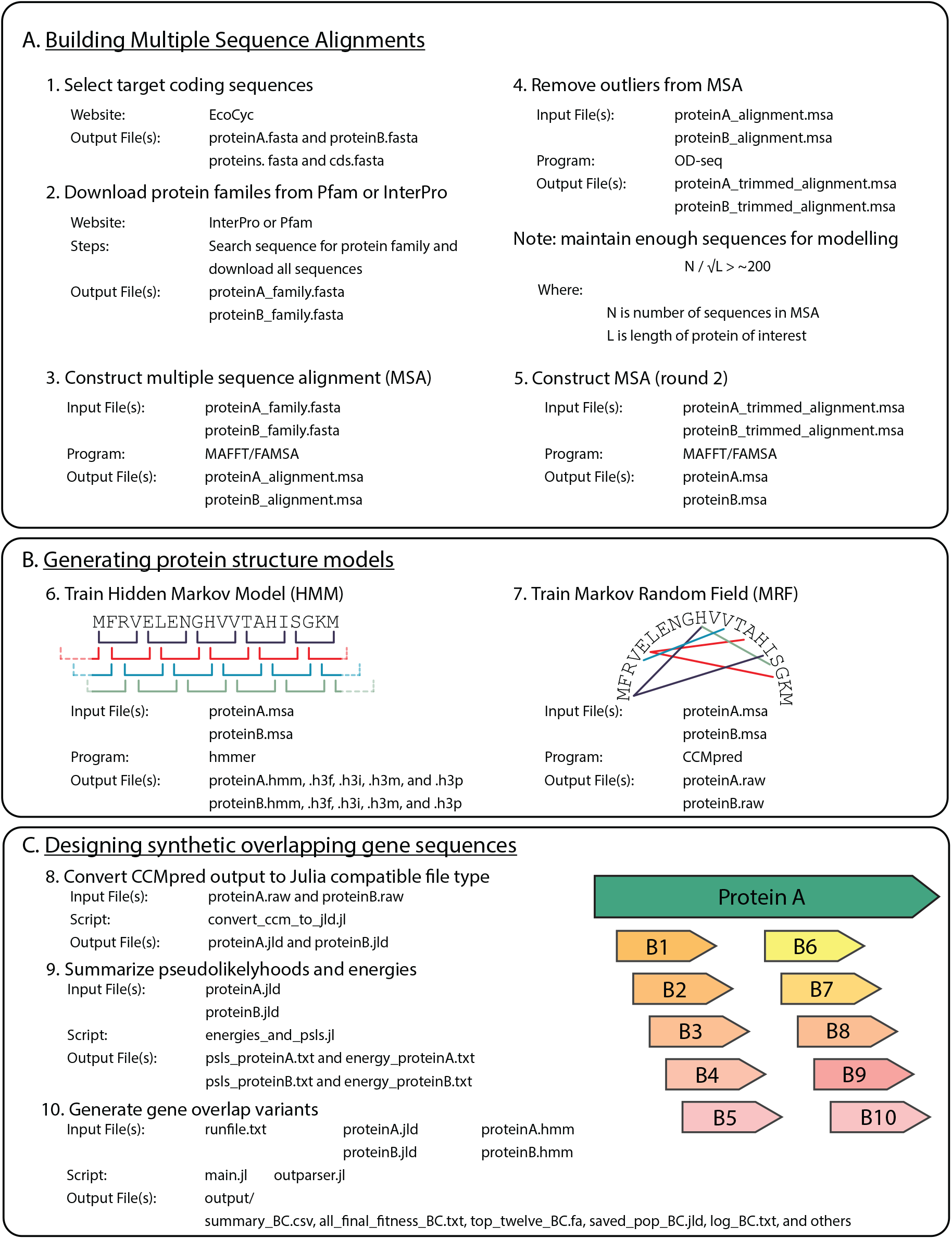
CAMEOS workflow. (A) The first step in the CAMEOS workflow is to create and curate MSA for the two target proteins. This process requires downloading protein family libraries from Pfam or InterPro, aligning these sequences through FAMSA or MAFFT, removing outliers via OD-seq, and repeating the alignment. (B) The second step is creating protein structure models (HMM and MRF) through hmmer and CMMpred. (C) The final step uses the CAMEOS scripts to generate the synthetic overlapping proteins and the library of the overlapping coding sequences.

Following the creation of an MSA of the two proteins to be co-encoded, HMMs and MRFs are generated (Figure 1B). A HMM operates by searching for patterns in a sequence space and calculates the probability of a pattern, or state, occurring (e.g. G, C, A, and T having a 25 % chance) and the transition probability of changing states (e.g. 75 % change of moving from state 1 to state 2). The hidden component is the transitions between states inside an observed sequence. The role of the HMM is to represent protein sequence conservation across protein family members [25,26]. A MRF is an undirected graphical probability model and represents combinations of independent assumptions which more directed models, such as Bayesian modelling, cannot accurately depict [25,26]. The role of the MRF is to represent intra-protein residue-residue coupling which may be crucial to protein function.

As the HMM detects conserved direct relationships and the MRF detects conserved indirect relationships, these models together create a fuller picture crucial of protein sequence and structure in targeted proteins [26]. HHMs are used in CAMEOS to create co-encoding solutions that are subsequently used as seeds in a second step where long-range interactions between protein residues are assessed with the MRFs [20].

In this chapter, we describe the steps needed to use CAMEOS to design *de novo* overlapped genes. We detail the processes to assemble sequences, and generate multiple sequence alignments, create HMMs and MRFs, run scripts included in the CAMEOS directory to pre-process the input data into the correct formats, and finally, to run the CAMEOS algorithm itself (Figure 1C).

## 2 Materials

### 2.1 Hardware

Intel Core i7-4770 3.40GHz with 4 cores and 32 GB RAM

### 2.2 Software

1. Ubuntu v20.10: https://ubuntu.com/download/desktop
2. hh-suite (v3.3.0) [27], an open-source package for sensitive protein sequence searching based on the pairwise alignment of Hidden Markov models (HMMs). GitHub: https://github.com/soedinglab/hh-suite
3. GCC v4.4+, a C compiler written for the GNU operating system. Website: https://gcc.gnu.org/
4. CMake v2.8+, an open-source cross-platform tool family to build, test, and package software. Website: https://cmake.org/
5. CCMpred [28], an open-source package for learning protein residue-residue contacts for building Markov Random Fields (MRF). GitHub: https://github.com/soedinglab/CCMpred
6. CAMEOS [20], an open-source package to generate *de novo* overlapped sequences. GitHub: https://github.com/BiosecSFA/CAMEOS
7. Julia (v1.4.1), a dynamic language for technical computing. With packages: BioAlignments, BioSymbols, Logging, StatsBase, JLD, Distributions, ArgParse, and NPZ.
8. Python (v3.9.5), an open-source cross-platform programming language. Website: https://www.python.org/
9. HDF5 (v1.10.6), a data software library and file format to manage, process, and store heterologous data. Website: https://www.hdfgroup.org/solutions/hdf5/
10. gzip (v1.10), a data compression program for the GNU operating system. Website: https://www.gnu.org/software/gzip/
11. hmmer v3+ [29], an open-source package for searching biological sequence databases for homologous sequences. GitHub: https://github.com/EddyRivasLab/hmmer
12. zlib1g-dev (v1.2.11) and groovy, a compression deflation method found in gzip and PKZIP, Website: https://packages.ubuntu.com/bionic/zlib1g-dev
13. OD-seq, an MSA analysis software package which detects outlier sequences. Download: http://www.bioinf.ucd.ie/download/od-seq.tar.gz
14. FASTX-Toolkit (v0.0.14), a collections of command line stools for Short-Reads FASTA/FASTQ files preprocessing. GitHub: https://github.com/agordon/fastx_toolkit
15. MAFFT (v7.310), a multiple sequence alignment program for Unix-like operating systems. Website: https://mafft.cbrc.jp/alignment/software/ https://anaconda.org/bioconda/mafft
16. FAMSA (v1.6.2), an algorithm for large-scale multiple sequence alignments. Website: https://github.com/refresh-bio/FAMSA https://anaconda.org/bioconda/famsa

## 3 Methods

Here, we describe the overall workflow to go from a pair of proteins we want to overlap to the output of DNA sequences that can be synthesized and tested in a wet lab. Complementary information to what is presented here can be found in the excellent `manual.pdf` file within the original CAMEOS GitHub repository (https://github.com/wanglabcumc/CAMEOS/tree/master/doc). Throughout this Methods section we use code that was downloaded from a fork of the original CAMEOS GitHub repository on 1 Dec 2021 (https://github.com/BiosecSFA/CAMEOS) that improved the original code in several ways. For details see notes (https://github.com/wanglabcumc/CAMEOS/pull/2). For a more comprehensive description of the development and theoretical underpinnings of the CAMEOS method, please see the original publication by Blazejewski et al. [20].

### 3.1 Choose protein sequences to overlap

The choice of which two proteins to overlap is nearly limitless, but there will be constraints based on sequence similarity and compatibility at the amino acid and DNA (coding sequence) level. The CAMEOS method was originally used to generate two sets of *E. coli* sequence pairs (CysJ-InfA and IlvA-CcdB), via >7,500 designs. A subset of these designs that were experimentally characterised showed that protein function and activity was maintained in both co-encoded proteins across their designs [20]. Additionally, 5.8 million theoretical overlaps between 199 essential genes and 49 non-essential biosynthetic gene sequences were computed. These analyses showed that 9 % of their computationally analysed subset contained pseudo-likelihood scores exceeding the experimentally characterised sequence pairs. From this, it was inferred that 80 % of the biosynthetic genes could be encoded with at least one essential gene. In this chapter we will use the *infA* (translation initiation factor IF-1) and *aroB* (3-dehydroquinate synthase) *E. coli* coding sequences (Notes 1 and 2) originally included as examples with the CAMEOS code on GitHub. All following examples will be just for InfA but it should be assumed that, where appropriate, the same process must also be done for AroB sequences. Additionally, because the AroB protein is longer, the analyses of this protein will take longer and may require more computational resources.

### 3.2 Download target protein and coding sequences

There are several sources of very large multiple sequence alignments (MSAs) that can be used as a starting point for a CAMEOS experiment. We will focus here on Pfam [30] and InterPro [31] databases, which are both large collections of protein families created and hosted by EMBL-EBI. We will use these databases as sources of large numbers of homologous sequences we can use to produce our own high-quality MSAs that are then fed into the CAMEOS workflow.

1. To determine how many protein sequences at minimum we will need for our alignments we can approximate using this formula: N/sqrt(L) > ∼200 Where N is number of sequences in MSA, sqrt() is the square root, and L is the length of protein of interest in amino acids (https://github.com/wanglabcumc/CAMEOS). For InfA with a length of 72 aa, the minimum number of sequences in the MSA would need to be N = sqrt(72) x 200 ≥ 1,697. For AroB, with a length of 362 amino acids, the minimum number would be more than double at 3,805 sequences. This number of sequences could be fulfilled from either Pfam or InterPro, but we will detail how to download sequences from InterPro as it provides a higher number.
2. First navigate to: https://www.ebi.ac.uk/interpro/search/text/
3. Keyword search for ‘IF-1’ since this is the protein encoded by *infA* gene (Note 3).
4. Potentially more accurate searching can also be done using the amino acid sequence of the protein of interest. In this case you would use the ‘Search -> By Sequence’ menu option of InterPro and enter the FASTA sequence of the protein you were interested in identifying the protein family of (Figure 2A).
5. Click on ‘ACCESSION’ link (IPR004368) for ‘Translation initiation factor IF-1’ from InterPro under the ‘SOURCE DATABASE’ heading.
6. Click on Proteins (46K) header, within this tab. Further filtering of the family can be performed to separate “reviewed” and “unreviewed” sequences. However, as the 634 reviewed proteins falls beneath the > 1,697 sequences required, we will continue with the unfiltered data.
7. Click on triangle portion of blue Export button on right hand side of page.
8. Hover the cursor over the Generate button beside the FASTA entry and you will see a popup window with ‘See more download options’ (Figure 2B). Click the button.
9. On the new page that opens, the ‘Choose a main data type’ header should be ‘Protein’
10. Scroll down and under ‘Select Output Format’ heading change to ‘FASTA’
11. Scroll down to bottom of page and click the Generate button (Notes 4 and 5).
12. When the data is ready for download the ‘Download’ button will light up (Figure 2C). Click this button and name the file `infA.fasta`

**Figure 2.**
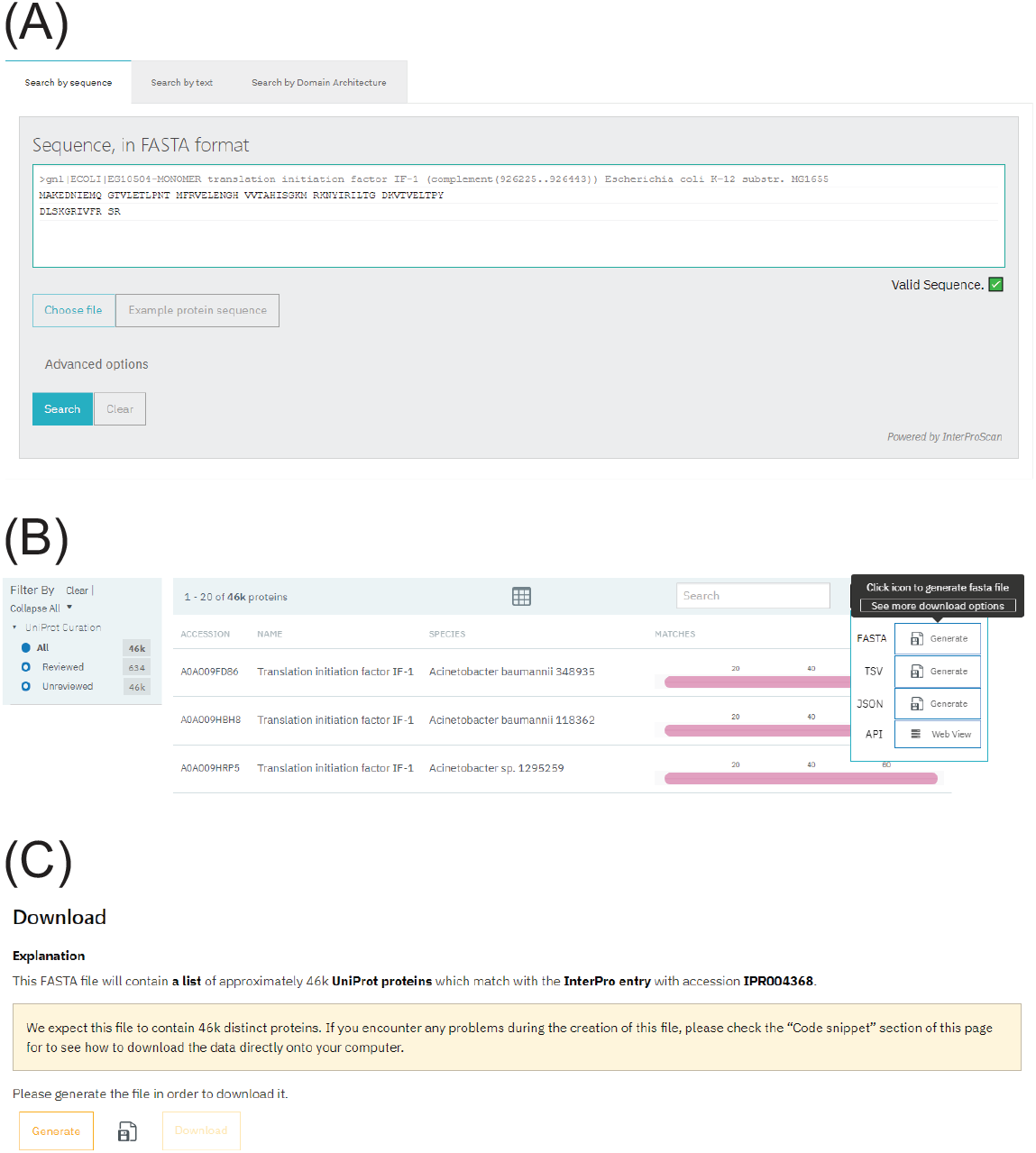
InterPro website navigation. (A) Search using the protein sequence of choice in the InterPro search bar to identify the protein family of the target protein. (B) After selecting the search results and navigating to the protein family, move to the Protein tab on the webpage, find the Export function, and click on the See More Download Options when hovering over the FASTA Generate button. (C) On this page, select the chosen data outputs and click Download.

### 3.3 Gathering additional sequences with HHBlits

While InterPro and PFam are excellent resources for downloading the sequences of protein families, sometimes more sequences are required to train the protein models than are provided by these sources. A way of gathering additional sequences is using the HHblits tool within HH-suite which iteratively searches sequences to detect similarities building high quality MSAs [27]. There are two methods for using HHblits for gathering aligned sequences: either using HHblits command `hhblits` on a CLI, or via the HHblits webtool.

The HHblits webtool (https://toolkit.tuebingen.mpg.de/tools/hhblits) will be discussed first as it is the simpler approach, although it offers fewer user input options. The tool requires a single protein sequence of interest or a MSA as the starting point. Additionally, the user is able to specify the following search parameters: (1) the Expect (E) value cut-off for inclusion, the (2) number of HHblits search iterations performed, the (3) minimum probability in the hitlist, and the (4) maximum number of target hits. All modification options are available within a dropdown menu and the default settings are clearly noted. Within a few minutes of submission, HHblits will return results listing the: (1) number of hits, (2) their identity, and (3) the alignment of those sequences (Figure 3A). For the downstream processes, the user must navigate to the “Query Template MSA” tab and download the full MSA file by clicking the option “Download Full MSA”. The file generated from this is in a *protein.a3m* file format and can be easily used with MAFFT without conversion to an protein.msa file extension (Figure 3B).

**Figure 3.**
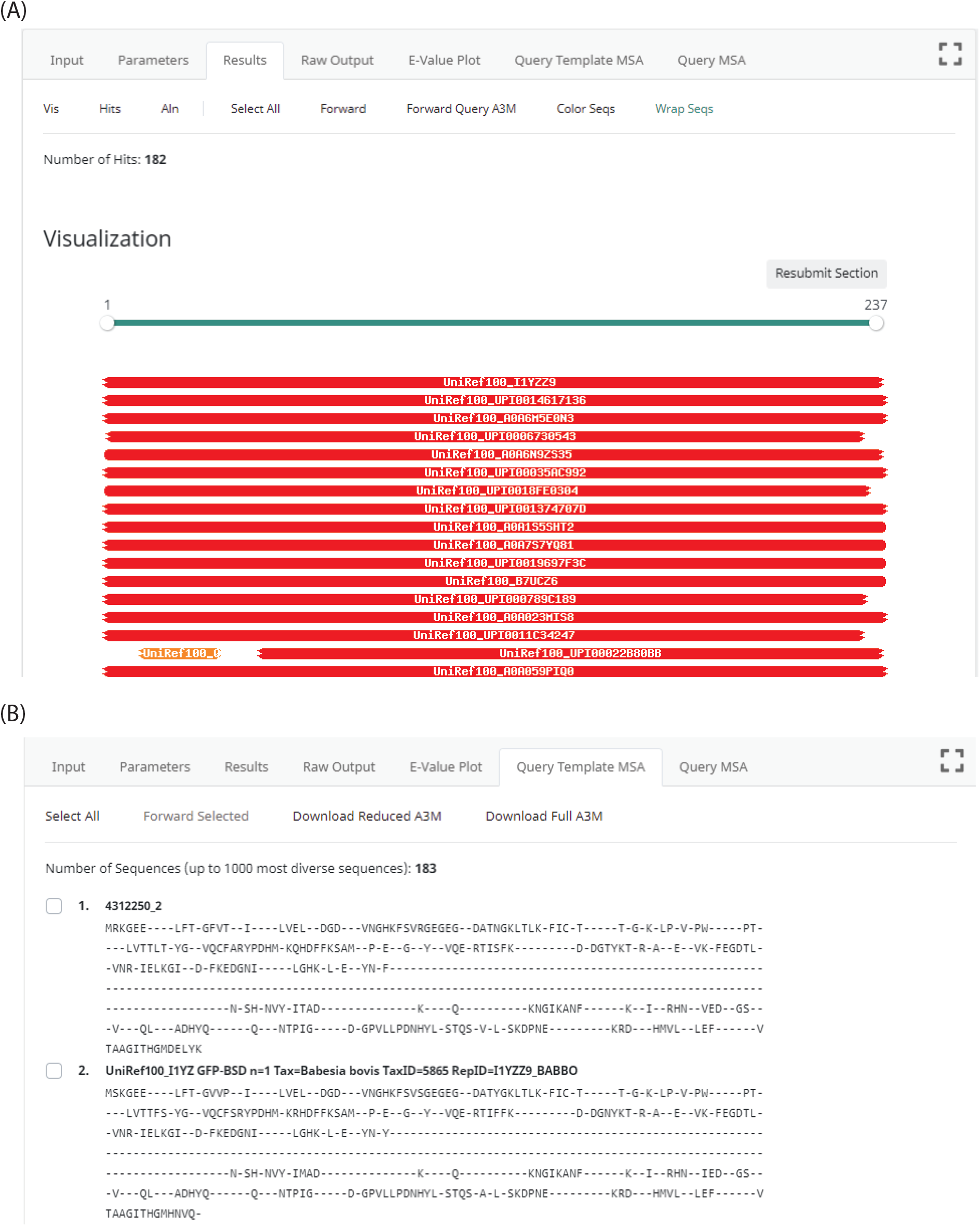
HHblits webserver outputs. (A) Once HHblits has queried the user input sequence, the server generates a visualisation output aligning returned sequences to the input sequence within the Results tab. (B) MSA (in the .a3m format) for the sequence search are accessed via the Query Template MSA tab and can be downloaded in Reduced or Full formats.

The other option to gather more sequences using the HHblits algorithm is to download the hh-suite package from GitHub directly (Note 6) and run `hhblits` from a CLI. The associated manual on `hhsuite` is very detailed and easy to use, however the tool requires a large downloaded sequence database (50+ GB) to function.

### 3.4 Gathering additional sequences with PSI-BLAST

An complementary approach to the protein domain focused databases InterPro and Pfam, and the search tool HHblits, is the algorithm Position-Specific Iterative BLAST, or PSI-BLAST. PSI-BLAST is a publicly available database search tool hosted by NCBI which performs an iterative search function against a protein query. Detailed instructions for PSI-BLAST are on the NCBI website and in this reference [32]. PSI-BLAST has some advantages over the previously mentioned tools as it provides increased sequence coverage by trading off poorer identity coverage. This is important as building the HMM will require a low number of gaps to generate a HMM profile that is usable with CAMEOS.

PSI-BLAST is an easy to use tool requiring a single protein sequence input. To use the PSI-BLAST function, users navigate to the protein BLAST (blastp) on NCBI, enter the input protein in the Query Sequence section, and select the PSI-BLAST algorithm in the Program Selection section. From the original input PSI-BLAST will generate search results matching the input protein limited by either sequence counts (default = 500), or by E value cut off (default is 0.005). These settings can be modified in the algorithm parameters on the initial search page to expand or constrict the available results from the first iteration. The initial results are used to seed the second iteration which is controlled by selecting the number of sequences to add in the “Run PSI-BLAST iteration 2” input. Search results can be filtered after by Percentage Identity, E value, Query coverage, and threshold cut offs. Further iterations can be performed to expand the sequence counts. All results can be downloaded from the browser as either aligned or unaligned sequences in FASTA format.

### 3.5 Perform multiple sequence alignment using MAFFT

The original CAMEOS publication used the FAMSA aligner [23] in concert with OD-seq [33] to remove outliers >2 standard deviations away from the sequences in a dataset, followed by manual removal of alignment positions when less than 50% of entries were aligned amino acids. Alternatively, we present another method below using the MAFFT aligner that seems to produce comparable results with less manual intervention.

1. Because the MAFFT aligner is not as efficient as FAMSA with large numbers of sequences (>10,000) it may be necessary to take a subsample of the files obtained from InterPro. The `fasta-subsample` tool in the MEME Suite [34] is an easy way to do this. Because of interference between different tools used in this protocol we used the Conda environment and package manager [35] to create a new environment just to run the MEME suite. All tools we use in this protocol were within the Ubuntu Linux operating system. After starting up a CLI: ~~~
(base) $ conda activate meme
~~~ Then from within that Conda environment where MEME suite has been installed you can use the `fasta-subsample` tool. Navigate to the folder containing the `infA.fasta` file, then using the CLI: `(meme) $ fasta-subsample infA.fasta 10000 >infA_sub_10000.fasta` Where: 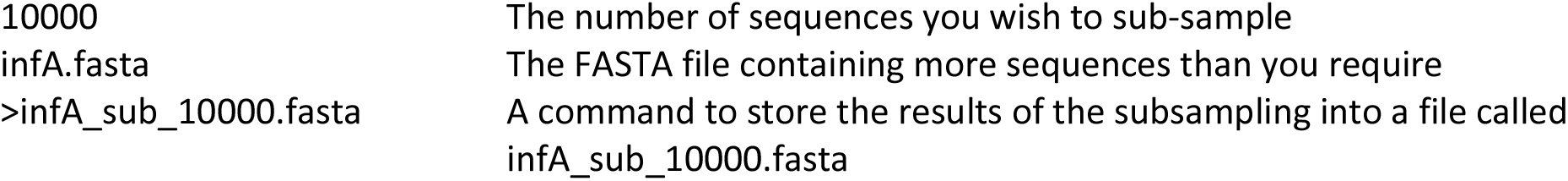 You can check that there are actually 10,000 FASTA files within this newly created file using the `grep` command and pipe the results to the `wc` command: ~~~
(meme) $ grep -o ‘>‘ infA_sub_10000.fasta | wc -l
10000
~~~
2. Running the MAFFT tool within our (base) environment on the newly created sub-sampled FASTA file of InfA sequences. ~~~
(base) $ time mafft --add infA_sub_10000.fasta --keeplength infA.fasta
>infA.msa
~~~ We use the `time` command to show us how long the alignment process took after it has completed. The `--add` flag is used to adding unaligned full-length sequence(s) into an existing alignment. In this case we are not adding to an alignment but using the sub-sampled InfA sequences in the file `infA_sub_10000.fasta`. This is done so that the alignment does not contain too many gaps and is relative to the target sequence. The `--keeplength` flag is used to chop off the ends of alignments that go over the *E. coli* InfA target sequence (`infA.fasta`), which simulates the effects of manual pruning (Note 7).
3. The sequence alignment may contain outlier sequences that would reduce the accuracy of the CAMEOS designs. To remove outlier sequences from the MSA we will use OD-seq [33] on the alignment (Note 8). ~~~
$ OD-seq -s 2 -i infA.msa -c infA_trim.msa
~~~ Where:

4. The MAFFT alignment is then repeated to align the sequences that were not removed by OD-seq.
5. The FASTA formatted output of MAFFT (and FAMSA) is not directly compatible with the CAMEOS scripts so must be converted to a single-line FASTA format. For CAMEOS to recognize the MSA files, each sequence, including gap characters (-), must occupy only one line (Note 9). We will use the `fasta_formatter` command of FASTX-Toolkit (Note 10) to do this: ~~~
$ fasta_formatter -i infA_trim.msa -o infA.msa -w 0
~~~

|  |  |
| --- | --- |
| OD-seq | The program that removes outlier sequences |
| -s | Flag to specify number standard deviations or greater from the mean the sequences need to be to remove from the alignment (in this case two standard deviations is selected). |
| -i | Flag to specify input file name (infA.msa) |
| -c | Flag to specify the output file name for sequences with average distance of less than two standard deviations to the rest of the sequences in the alignment (infA_trim.msa) |
| -o | (optional) Flag to specify the output file name for sequences with average distance of more than two standard deviations to the rest of the sequences in the alignment. |

Where:

|  |  |
| --- | --- |
| fasta_formatter | Tool used to reformat FASTA sequences |
| -i | Flag to specify input file name (infA_trim.msa) |
| -o | Flag to specify output file name (infA.msa) |
| -w | Flag to specify the max. sequence line width for output FASTA file. The 0 means that sequences lines will not be wrapped and all amino acids will appear on the same line. |

### 3.6 Creation of a Protein Generative Model

Before two protein sequences can be artificially overlapped with the CAMEOS algorithm, each protein sequence must be analysed to create both a HMM and a MRF representation. This is done so that the CAMEOS algorithm can determine regions of the proteins where sequence flexibility and long-range interactions (residue-residue contacts) are amenable to coding sequence overlap in different reading frames. Unless otherwise stated, we assume the `proteinA.msa` and `proteinB.msa` files (in our case `infA.msa` and `aroB.msa`) are within the `main/` sub-folder of the CAMEOS script folder and your CLI program’s present working directory (`pwd`) is also `main/`

#### 3.6.1 Training HMM using hmmer

1. Using our generated `infA.msa` and `aroB.msa` files we will first generate hidden Markov models (HMMs) of each protein using hmmer (http://hmmer.org/) command `hmmbuild`. ~~~
$ hmmbuild infA.hmm infA.msa
~~~ Where:

2. Inspect the resulting information generated in the CLI to determine if the HMM was generated correctly. The CAMEOS script is able to use .hmm files that are incomplete without raising an error, but the end results of the process will be incorrect. An indication that your HMM files are not correct is the final engineered protein sequences generated will include gaps and will be shorter than the actual protein sequences put into the script in the `proteins.fasta` file. To ensure these errors do not occur, your HMM must be of the <u>same length as the input protein</u>. In the `hmmbuild` output check that ‘alen’ and ‘mlen’ are the same value (72 in this case) and that this value is the same as the length of the protein in amino acids (72 aa in this case is the full length of InfA).

|  |  |
| --- | --- |
| `hmmbuild` | The function of hmmer that builds a HMM |
| `infA.hmm` | Output file name |
| `infA.msa` | Input file name |

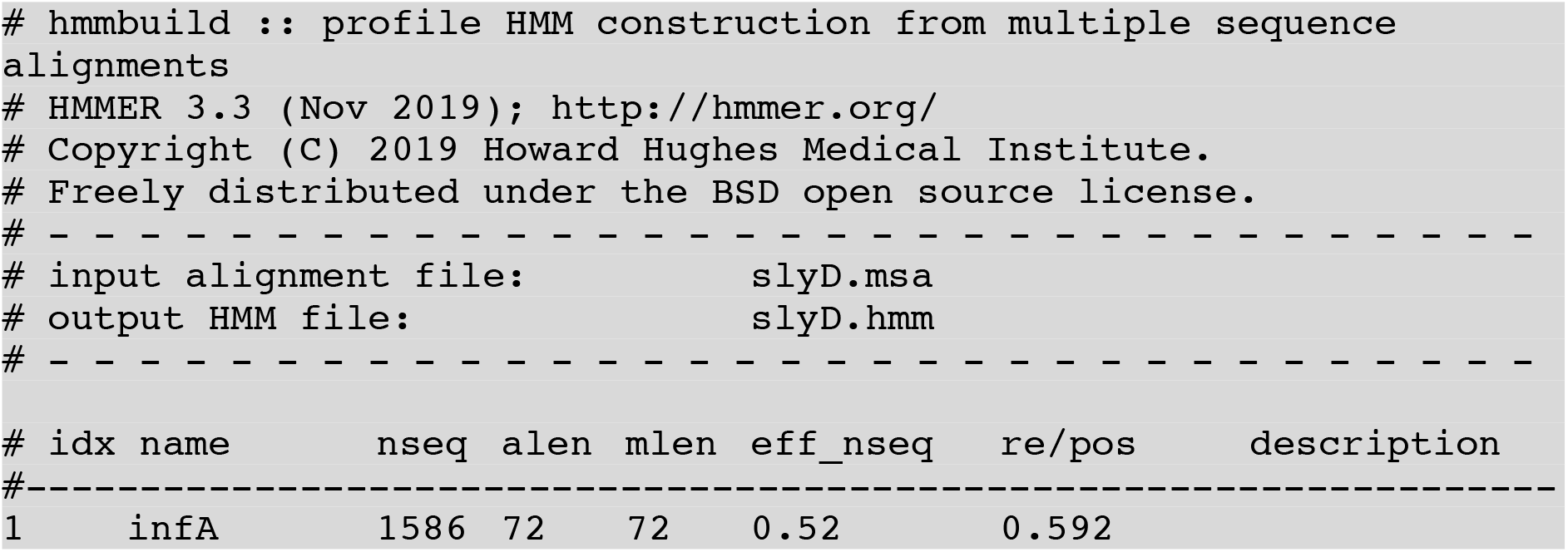
3. The generated .hmm file is then compressed to create the final .hmm/.h3f/.h3i/.h3m/.h3p files CAMEOS requires, using the `hmmpress` command of the `hmmer` package.

~~~
$ hmmpress infA.hmm
~~~

Where:

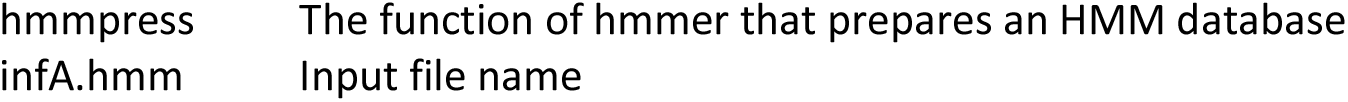

#### 3.6.2 Training Markov Random Field using CCMpred

The MRF model for each protein will be trained using CCMpred [28] to create residue-residue contact predictions. These models are for later use in assessing the impact of protein sequence changes and their long-range interactions within a protein family.

1. First we must convert the MSA files to a format that is compatible with CCMpred (only sequences, no FASTA headers) using an inverse-match `grep` command:

~~~
$ grep -v “>“infA.msa > infA.ccm
~~~ Where:

2. Next we invoke CCMpred to generate a .raw matrix file.

|  |  |
| --- | --- |
| `grep` | A command-line utility for searching lines that match a regular expression. |
| `-v` | Inverse-match flag |
| `">"` | The string to match in the input file |
| `infA.msa` | Input file |
| `>` | Save results of `grep` to a file |
| `infA.ccm` | Output file name |

~~~
$ ccmpred -t 1 -r infA.raw -n 100 infA.ccm infA.mat
~~~ Where:

3. The .raw file is not in the correct format expected by the CAMEOS scripts, so it must be converted to an internal MRF file format using a Julia [36] language script (`convert_ccm_to_jld.jl`). The script generates a Julia Data File (.jld) file that is used when the `main.jl` CAMEOS Julia script is run later:

|  |  |
| --- | --- |
| `ccmpred` | The ccmpred command |
| `-t 1` | (optional) Depending on the number of CPUs you have available for the computation, you may want to use a value >1 (default) here to complete the calculation faster |
| `-r` | Store raw prediction matrix in RAWFILE format flag |
| `infA.raw` | The output raw file |
| `-n 100` | Compute a maximum of NUMITER operations [default: 50] |
| `infA.ccm` | The input file name |
| `infA.mat` | The matrix output file |

~~~
$ julia convert_ccm_to_jld.jl infA.raw infA.jld
~~~

Where:

|  |  |
| --- | --- |
| `julia` | Runs script with Julia |
| `convert_ccm_to_jld.jl` | Name of script to run |
| `infA.raw` | Input file name |
| `infA.jld` | Output file name |

The .jld files must then be transferred to the `jlds/` sub-folder or the `main.jl` script will fail when run later.

#### 3.6.3 Summarizing pseudolikelihoods/energies

1. Next we must summarize the data from CCMpred into formats that work with the CAMEOS scripts. The pseudolikelihoods and energies of the proteins are used in the optimization process and so these values are calculated using the `energies_and_psls.jl` script that will output two files: `psls_protein.txt` and `energy_protein.txt` into the `psls/` and `energies/` sub-folders, respectively. These folders must already exist within the `main/` folder or the `main.jl` script will fail when run. In our example the files would be named `psls_infA.txt` and `energy_infA.txt`

The script is run:

~~~
$ julia energies_and_psls.jl infA infA.jld infA.msa
~~~

Where:

|  |  |
| --- | --- |
| `julia` | Runs script with Julia |
| `energies_and_psls.jl` | Name of script to run |
| `infA` | The name of the protein |
| `infA.jld` | Input Julia Data File name. Note, this was generated in the previous step. |
| `infA.msa` | Input MSA file name. Note, this was generated in a previous step. |

#### 3.6.4 Setting up folder structure

The `main.jl` script requires all the input files to be in a certain folder structure or it will fail. Within the `main/` folder the correct subfolder structure is:

~~~
energies/
~~~

Containing: `energy_infA.txt` and `energy_aroB.txt` files.

~~~
hmms/
~~~

Containing: `infA.hmm, infA.hmm.h3f, infA.hmm.h3i, infA.hmm.h3m, infA.hmm.h3p, aroB.hmm, aroB.hmm.h3f, aroB.hmm.h3i, aroB.hmm.h3m, and aroB.hmm.h3p` files.

~~~
jlds/
~~~

Containing: `infA.jld` and `aroB.jld`

NOTE: as of the time of this writing (early 2022) GitHub does not host these files correctly and instead of `aroB.jld` being ∼464.9 MB and `infA.jld` being ∼18.3 MB, they instead are 134 bytes and 133 bytes, respectively. The reduced-size files will cause an error if used as-is. Two options to get around this limitation of GitHub are to download the correct files here: https://cloudstor.aarnet.edu.au/plus/s/jpM0fvly0Y2r4Wi

Alternatively, if you run the CAMEOS process from the beginning, as described in this chapter, you will generate your own `aroB.jld` and `infA.jld` files of the correct size.

~~~
msas/
~~~

Containing: `infA.msa` and `aroB.msa`

~~~
output/
~~~

Containing: nothing. The folder is empty at the start of the script.

~~~
psls/
~~~

Containing: `psls_infA.txt` and `psls_aroB.txt`

Additionally, within the `main/` folder the following files are required:

~~~
cds.fasta, proteins.fasta, runfile.txt
~~~

#### 3.6.5 Running CAMEOS

1. The last step before running the main CAMEOS script is to modify the file containing the run parameters. Here, we call the file `runfile.txt` but you can name it whatever makes sense to you. The file contains the parameters used during the `main.jl` script execution and controls aspects of the CAMEOS process, such as how many seeds to optimize and how many iterations to perform.

These parameters must be adjusted carefully because they can have dramatic effects, such as significantly extending runtime.

The `runfile.txt` parameter file has the following structure:

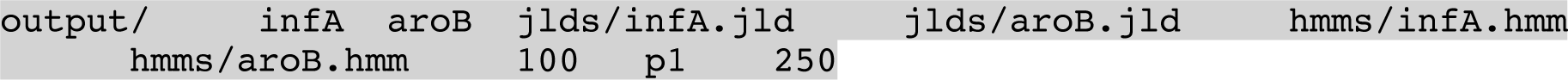

The file is tab-delimited (each unit of text is separated by a tab character) and stores the following information (Note 11), where:

|  |  |
| --- | --- |
| output/ | Directory where output files are stored |
| infA | Gene/Protein A name |
| aroB | Gene/Protein B name |
| jlds/infA.jld | Path to .jld file containing MRF (CCMpred data) for Gene A |
| jlds/aroB.jld | Path to .jld file containing MRF (CCMpred data) for Gene B |
| hmms/infA.hmm | Path to HMM file (hmmer data) for Gene A |
| hmms/aroB.hmm | Path to HMM file (hmmer data) for Gene B |
| 100 | Number of seeds to optimize |
| p1 | Frame. This parameter should not be modified |
| 250 | Number of iterations of algorithm |

The time the CAMEOS method takes to complete depends on a number of factors such as sequence lengths, seed value, and iteration value.

To run CAMEOS using the `main.jl` script, navigate to the `main/` folder and type:

~~~
$ julia main.jl runfile.txt
~~~

Where:

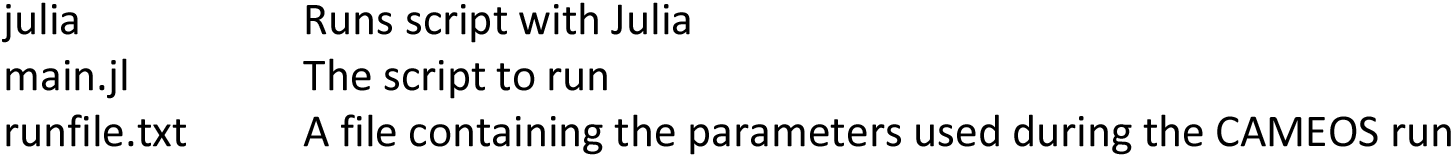

|  |  |
| --- | --- |
| julia | Runs script with Julia |
| main.jl | The script to run |
| runfile.txt | A file containing the parameters used during the CAMEOS run |

After a successful run, text will be displayed on the CLI, similar to:

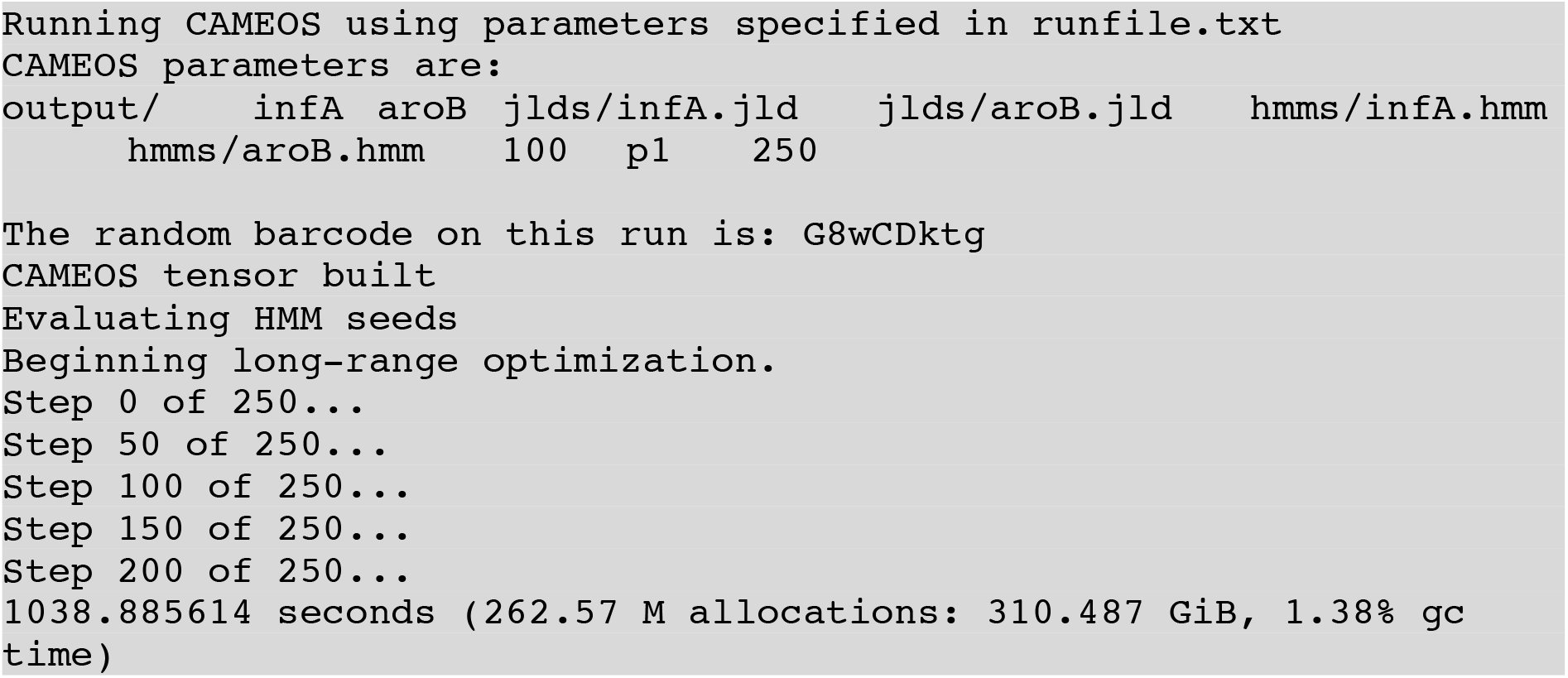

#### 3.6.6 Evaluating CAMEOS Results

1. In the `output/` subfolder, a number of files are created from a successful CAMEOS run. The `top_twelve_BC.fa` (where BC is barcode of the run. In our example above it would be `top_twelve_G8wCDktg.fa`) file contains the best three co-encodings of the two genes of interest from the best score of proteinA (InfA in our example) and proteinB (AroB in our example). Additionally, the file also contains co-encodings (CDS overlaps) with the best overall score. Although in most instances the script will just fail if the initial files are not of the correct type and location, we have seen a few cases where output is generated but is erroneous. For example, if the MSA files that are used have more characters in them than the sequences in the `proteins.fasta` file, the output sequences that are generated will have gaps in them. Therefore, careful analysis of the output sequences should be done before synthesizing the DNA to make the constructs.
2. An additional script (from: https://github.com/BiosecSFA/CAMEOS) can be used to summarize the information from the .jld output file into FASTA and comma separated values (CSV) file**s**, which are generally easier to look through. With the CLI’s present working directory as `main/`, execute the following code:

~~~
$ julia outparser.jl infA aroB BC --fasta
~~~ Where:

2. In the example of *infA* encoded within the *aroB* gene, we can see the top scoring hits are located in either the 5’ or 3’ regions of *aroB* (Figure 4A). Within the 5’ region, three InfA variant designs modified the AroB sequence on average 14% to enable the co-encoding of InfA into AroB. A similar result was observed in the 3’ region of *aroB* as the five InfA variants there modified AroB on average 13%.

|  |  |
| --- | --- |
| `julia` | Runs script with Julia |
| `outparser.jl` | Julia script that parses the CAMEOS output into easier to analyse files |
| `infA` | The name of the first protein co-encoded |
| `aroB` | The name of the second protein co-encoded |
| `BC` | Barcode from your CAMEOS run (e.g. G8wCDktg) |
| `--fasta` | Generates a FASTA file of the results in addition to a CSV |
| `--just-fullseq` | (optional): this flag can be used in addition to the `--fasta` flag to create a FASTA file that only contains the full sequence. |

**Figure 4.**
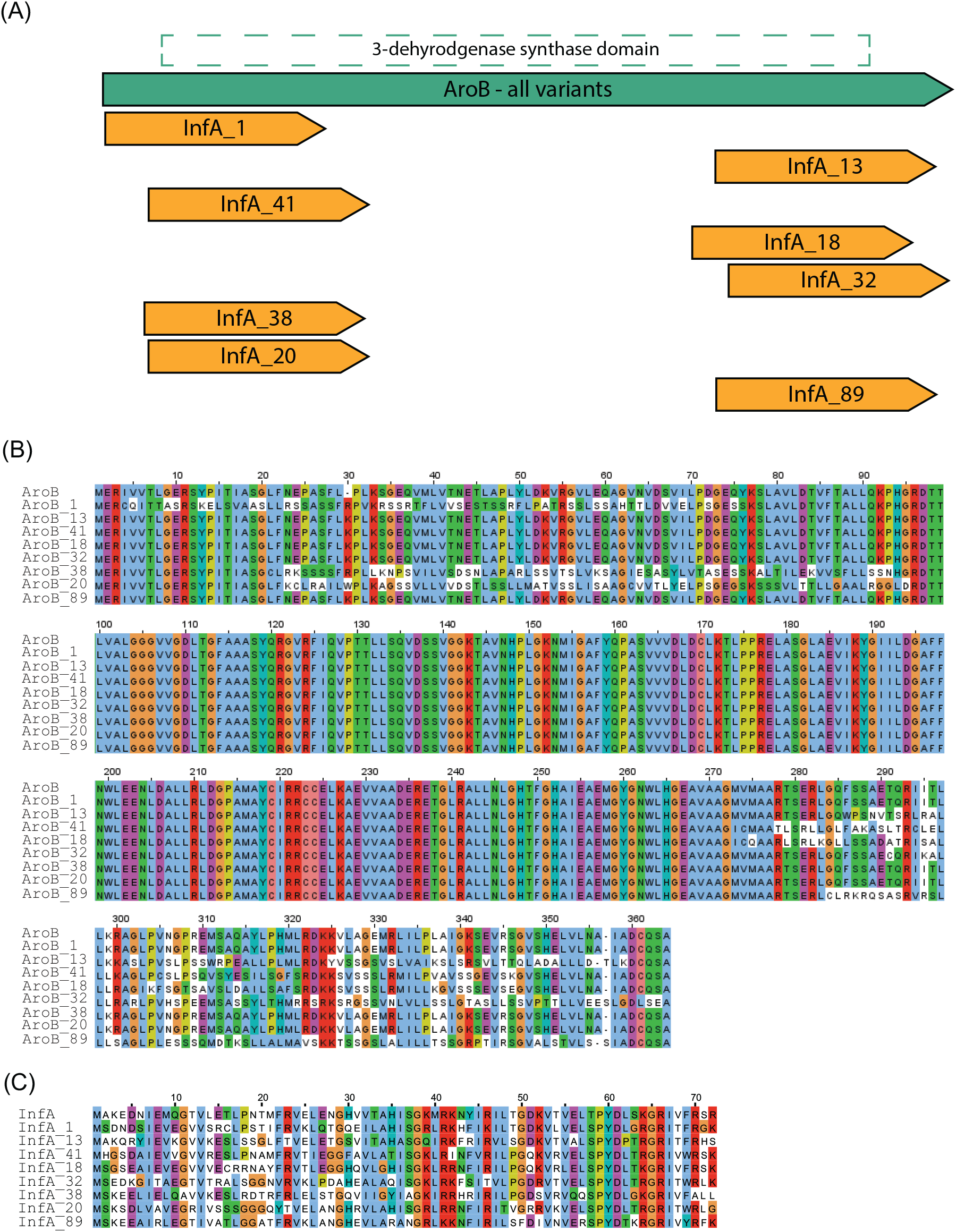
Aligned CAMEOS AroB and InfA outputs. (A) The eight highest scoring CAMEOS designs incorporated InfA within AroB within two regions, at the 5’ and 3’ ends. (B) AroB designs aligned to the wild-type AroB sequence showing the locations where residues were modified. (C) InfA designs aligned to the wild-type InfA sequence showing the locations where residues were modified. Despite being in two distinct regions, all designs incorporated a new residue at position 30. In most designs this was lysine, however in designs 1 and 20 an arginine and tryptophan were incorporated respectively (Figure 4B). While no crystal structure is available for *E. coli* AroB, on UniProt an Alphafold simulation is publicly available predicting the protein structure [37,38]. The inserted AroB residue is incorporated 5’ adjacent to a proline residue which terminates a predicted alpha helix. All three modified residues have some favourability to form alpha helices, therefore it is likely that the modification either continues the alpha helix one residue or has a secondary structural effect. Overall, AroB is predicted to be a highly structured protein, therefore all modified residues will interact with existing secondary structures. For example, in the 5’ region containing InfA designs the existing structure is a combination of alpha helices and beta sheets, while the 3’ region containing InfA designs is populated with predominantly alpha helices with a single beta sheet. Due to its short size, InfA had more significant modifications to its sequence as on average 45% of the amino acid identities were altered (Figure 4C). Unlike AroB, InfA has an experimentally characterised structure which is dominated by a beta barrel with a single short alpha helix [39]. Therefore, due to InfA’s small size and highly structured topology, all residue changes would be members of an existing beta sheet or alpha helix.
3. Differences in the co-encodings can also be seen when considering the predicted translation efficiencies of *infA* from within the *aroB* sequence. Using the RBS Calculator [40] on the top five designs we see a nearly 8-fold difference between predicted translation efficiency of the worst and best *infA* designs. Similarly, there is a 9-fold difference for *aroB* (Figure 5A). The correlation between *aroB* and *infA* translation initiation rates in this case is due to the N-terminal location of *infA* co-encoding. If *infA* is co-encoded more C-terminally, the two CDS translation initiation rates (TIRs) are not connected, with *aroB* displaying a strong TIR (5,577 AU) and *infA* displaying a range of TIRs (1 - 286 AU), although 20 - 5,577-fold lower than *aroB* (Figure 5B). For reference, in the *infA* natural *E. coli* genomic context, it has a predicted TIR of 1,211 (AU) which is at least 4-fold higher than the best CAMEOS encoding.

**Figure 5.**
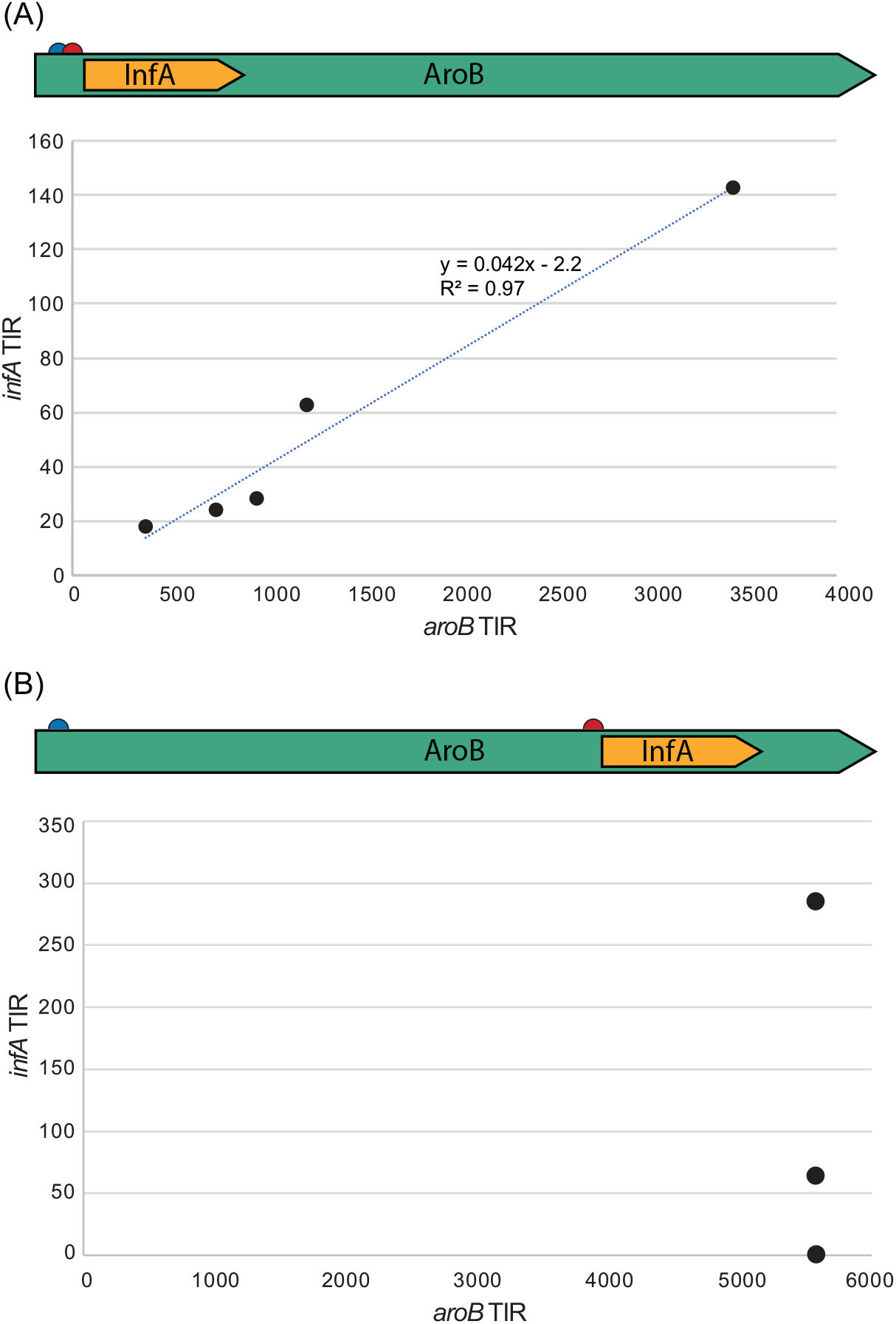
Predicted Translation Efficiencies for CAMEOS designs. (A) Using RBS Calculator on the top 5 N-terminal designs of *infA*/*aroB* overlap we see a strong correlation between *aroB* translation initiation rate (TIR) and the downstream *infA* TIR. This effect is likely due to the close proximity of the start codons and ribosome binding sites of the co-encoded coding sequences. (B) Using RBS Calculator on other *aroB*/*infA* co-encodings where the start codons and RBSs are spaced further apart shows no correlation between TIRs but does show as before, a wide range of *infA* TIRs.

### 3.7 Putting it all together

As we have outlined, the successful completion of co-encoding two proteins in the same DNA sequence in different reading frames, is a complex and multistep process. One of the most burdensome barriers to entry for molecular biologists is access to a computer running Linux and the installation of all the tools needed.

To ease this process and enable scientists without strong computational backgrounds to use the CAMEOS algorithm, we have created a virtual machine which comes pre-loaded with all the tools used in this protocol and can be easily run on a computer using Windows or macOS operating systems. The virtual machine file is ∼20GB large so ensure you have plenty of room on your disk and a fast internet connection. The disk size of the virtual machine is 50GB, so as you add data to your virtual machine ensure you have at least 75GB free on your computer’s disk.

Access the .ova file which contains the virtual machine (called ‘overlap’) here: https://cloudstor.aarnet.edu.au/plus/s/8ooJ7SHTt463KE5

The starting password for the Ubuntu operating system on the virtual machine is: overlap We strongly suggest you change the password once you start using the virtual machine.

To check that the `overlap.ova` file downloaded correctly on macOS, start up a CLI such as Terminal in the directory you downloaded the file to and run:

~~~
$ md5 overlap.ova
~~~

To check that the `overlap.ova` file downloaded correctly on Linux, start up a CLI such as Terminal in the directory you downloaded the file to and run:

~~~
$ md5sum overlap.ova
~~~

To check that the `overlap.ova` file downloaded correctly on Windows, start up a CLI such as Command Prompt in the directory you downloaded the file to and run:

~~~
C:\> certutil -hashfile overlap.ova MD5
~~~

The result of these commands should be: `b8f894df3305507bdee6e992ac87d75f`

If your result does not match, then it is likely that the download was interrupted and the `overlap.ova` file was corrupted. Please try to download again. In the future, if the link to this resource becomes broken, please check our lab website for details: https://www.jaschke-lab.science/

The virtual machine .ova file (`overlap.ova`) can be booted up using the free Oracle VM Virtualbox software.

2. We have also created a script in the shell language Bash that can be used to accomplish all the previously described steps automatically, reducing the chances of human error from moving all these files around and using tools with certain parameters. This Bash script is called `run_cameos.sh` and is stored within the `main/` folder of the CAMEOS code on the overlap.ova virtual machine. To just download the Bash script please find it here: https://cloudstor.aarnet.edu.au/plus/s/Q09KhQAQCaNimdj

To use the script to perform a run of CAMEOS, you need to open the `run_cameos.sh` script in a text editor, and specify the protein names you are working on by changing the two variables specifying the protein names:

~~~
proteinA=infA
protein=aroB
~~~

Then save and close the `run_cameos.sh` script file. Next, open a CLI in the `main/` folder and run the script by:

~~~
$ bash run_cameos.sh
~~~

As the script is running it will display updates on which tool is being run and its progress in the CLI window. Once the run is done it will display information on where the output files are located and how long the script took to run.

## 4 Notes

1. More information on these coding sequences can be seen on the Ecocyc database [41] here: https://ecocyc.org/gene?orgid=ECOLI&id=EG10504 https://biocyc.org/gene?orgid=ECOLI&id=EG10074
2. Only *E. coli* sequences have been used with CAMEOS before, although in principle nothing is preventing other prokaryote coding sequences from being used, any sequence differing from the standard codon table or *E. coli* codon usage would need to manually optimize the code.
3. Text to enter will be supplied with single quotes ‘text to be entered’ and the quotes should not be included unless specifically stated.
4. Depending on the number of sequences this process may take more than 1 hour to generate the data.
5. The FASTA sequences could also be downloaded programmatically using the available Application Programming Interface (API) using Python, Perl, or JavaScript. InterPro makes this process easier by automatically generating the code needed, but this method is outside the scope of the current article.
6. The HH-suite GitHub page has a detailed wiki page with examples of how to run their script and is accessible via https://github.com/soedinglab/hh-suite/wiki
7. https://mafft.cbrc.jp/alignment/server/add.html
8. Although available through Bioconductor for the R language, we used the CLI version available here from the original publication: http://www.bioinf.ucd.ie/download/od-seq.tar.gz
9. Some multiple sequence aligners (MAFFT and FAMSA, for example) create output with 50 or 80 characters per line separated by newline (\n) characters, which is not suitable for use in the CAMEOS script.
10. The FASTX-toolkit is available for download through both GitHub (https://github.com/agordon/fastx_toolkit) and through a website (http://hannonlab.cshl.edu/fastx_toolkit/).
11. The names used for proteins A and B must be identical to those used in the `proteins.fasta` and `cds.fasta` files in the FASTA header (e.g. >infA). The file names of the files storing the energies and pseudolikelihoods must also include the same identifier (e.g. `energy_infA.txt` and `psls_infA.txt`). Lastly, use a short identifier, such as the 3-4 letter gene name, because output files and folders will be named with these identifiers. For reference, the CAMEOS publication (Figure S2) showed the effects of using different values for Number of Iterations. Most of the tested variants did not display dramatically improved summed pseudolikelihood scores after ∼300 iterations. The Frame value p1 should not be altered as it is the only option currently supported.

## Acknowledgement

We recognize that the intellectual and physical labour of this research was conducted on the traditional lands of the Wallumattagal clan of the Dharug nation, the Gadigal, Wangal, and Cameraygal peoples of the Eora nation. We thank G Sullivan for assistance with FASTX-toolkit; T Blazejewski for explaining in detail to us how CAMEOS works; JM Marti, J Allen, Y Jiao, D Park, and D Ricci for helpful discussions and their improvements to the original CAMEOS code. DYL was supported by the Macquarie University COVID Recovery Postdoctoral Fellowship. PRJ was supported by NHMRC Ideas Grant APP1185399.

## References

1. Tang T-C, An B, Huang Y, Vasikaran S, Wang Y, Jiang X, Lu TK, Zhong C (2021) Materials design by synthetic biology. Nature Reviews Materials 6 (4):332–350. doi:10.1038/s41578-020-00265-w

2. Brophy JAN, Voigt CA (2014) Principles of genetic circuit design. Nature Methods 11 (5):508–520. doi:10.1038/nmeth.2926

3. Chan LY, Kosuri S, Endy D (2005) Refactoring bacteriophage T7. Molecular systems biology 1 (1):2005.0018

4. Jaschke PR, Lieberman EK, Rodriguez J, Sierra A, Endy D (2012) A fully decompressed synthetic bacteriophage øX174 genome assembled and archived in yeast. Virology 434 (2):278–284

5. Logel DY, Trofimova E, Jaschke PR (2022) Codon-Restrained Method for Both Eliminating and Creating Intragenic Bacterial Promoters. ACS Synthetic Biology. doi:10.1021/acssynbio.1c00359

6. Temme K, Zhao D, Voigt CA (2012) Refactoring the nitrogen fixation gene cluster from Klebsiella oxytoca. Proceedings of the National Academy of Sciences 109 (18):7085–7090

7. Wright BW, Molloy MP, Jaschke PR (2021) Overlapping genes in natural and engineered genomes. Nature Reviews Genetics. doi:10.1038/s41576-021-00417-w

8. Song M, Sukovich DJ, Ciccarelli L, Mayr J, Fernandez-Rodriguez J, Mirsky EA, Tucker AC, Gordon DB, Marlovits TC, Voigt CA (2017) Control of type III protein secretion using a minimal genetic system. Nat Commun 8:14737. doi:10.1038/ncomms14737

9. Springman R, Molineux IJ, Duong C, Bull RJ, Bull JJ (2012) Evolutionary stability of a refactored phage genome. ACS Synth Biol 1 (9):425–430. doi:10.1021/sb300040v

10. Borkowski O, Ceroni F, Stan G-B, Ellis T (2016) Overloaded and stressed: whole-cell considerations for bacterial synthetic biology. Current opinion in microbiology 33:123–130

11. Ellis T (2019) Predicting how evolution will beat us. Microbial biotechnology 12 (1):41

12. Gallup O, Ming H, Ellis T (2021) Ten future challenges for synthetic biology. Engineering Biology 5 (3):51–59. doi:https://doi.org/10.1049/enb2.12011

13. Clark M, Maselko M (2020) Transgene Biocontainment Strategies for Molecular Farming. Frontiers in Plant Science 11. doi:10.3389/fpls.2020.00210

14. Maselko M, Heinsch SC, Chacón JM, Harcombe WR, Smanski MJ (2017) Engineering species-like barriers to sexual reproduction. Nature Communications 8 (1):883. doi:10.1038/s41467-017-01007-3

15. Ryu M-H, Zhang J, Toth T, Khokhani D, Geddes BA, Mus F, Garcia-Costas A, Peters JW, Poole PS, Ané J-M, Voigt CA (2020) Control of nitrogen fixation in bacteria that associate with cereals. Nature Microbiology 5 (2):314–330. doi:10.1038/s41564-019-0631-2

16. Rylott EL, Bruce NC (2020) How synthetic biology can help bioremediation. Current Opinion in Chemical Biology 58:86–95. doi:https://doi.org/10.1016/j.cbpa.2020.07.004

17. Voigt CA (2020) Synthetic biology 2020–2030: six commercially-available products that are changing our world. Nature Communications 11 (1):6379. doi:10.1038/s41467-020-20122-2

18. Jaschke PR, Dotson GA, Hung KS, Liu D, Endy D (2019) Definitive demonstration by synthesis of genome annotation completeness. Proceedings of the National Academy of Sciences 116 (48):24206–24213

19. Wright BW, Ruan J, Molloy MP, Jaschke PR (2020) Genome modularization reveals overlapped gene topology is necessary for efficient viral reproduction. ACS Synthetic Biology 9 (11):3079–3090

20. Blazejewski T, Ho H-I, Wang HH (2019) Synthetic sequence entanglement augments stability and containment of genetic information in cells. Science 365 (6453):595–598. doi:doi:10.1126/science.aav5477

21. Chowdhury B, Garai G (2017) A review on multiple sequence alignment from the perspective of genetic algorithm. Genomics 109 (5):419–431. doi:https://doi.org/10.1016/j.ygeno.2017.06.007

22. Thompson JD, Higgins DG, Gibson TJ (1994) CLUSTAL W: improving the sensitivity of progressive multiple sequence alignment through sequence weighting, position-specific gap penalties and weight matrix choice. Nucleic Acids Res 22 (22):4673–4680

23. Deorowicz S, Debudaj-Grabysz A, Gudyś A (2016) FAMSA: Fast and accurate multiple sequence alignment of huge protein families. Scientific Reports 6 (1):33964. doi:10.1038/srep33964

24. Katoh K, Misawa K, Kuma Ki, Miyata T (2002) MAFFT: a novel method for rapid multiple sequence alignment based on fast Fourier transform. Nucleic Acids Res 30 (14):3059–3066. doi:10.1093/nar/gkf436

25. Eddy SR (2004) What is a hidden Markov model? Nature Biotechnology 22 (10):1315–1316. doi:10.1038/nbt1004-1315

26. Thomas J, Ramakrishnan N, Bailey-Kellogg C (2008) Graphical models of residue coupling in protein families. IEEE/ACM Transactions on Computational Biology and Bioinformatics 5 (2):183–197

27. Steinegger M, Meier M, Mirdita M, Vöhringer H, Haunsberger SJ, Söding J (2019) HH-suite3 for fast remote homology detection and deep protein annotation. BMC Bioinformatics 20 (1):473. doi:10.1186/s12859-019-3019-7

28. Seemayer S, Gruber M, Söding J (2014) CCMpred--fast and precise prediction of protein residue-residue contacts from correlated mutations. Bioinformatics 30 (21):3128–3130. doi:10.1093/bioinformatics/btu500

29. Finn RD, Clements J, Eddy SR (2011) HMMER web server: interactive sequence similarity searching. Nucleic Acids Res 39 (Web Server issue):W29–W37. doi:10.1093/nar/gkr367

30. Finn RD, Bateman A, Clements J, Coggill P, Eberhardt RY, Eddy SR, Heger A, Hetherington K, Holm L, Mistry J (2014) Pfam: the protein families database. Nucleic Acids Res 42 (D1):D222–D230

31. Blum M, Chang H-Y, Chuguransky S, Grego T, Kandasaamy S, Mitchell A, Nuka G, Paysan-Lafosse T, Qureshi M, Raj S (2021) The InterPro protein families and domains database: 20 years on. Nucleic Acids Res 49 (D1):D344–D354

32. Bhagwat M, Aravind L (2007) PSI-BLAST Tutorial. In: Nh B (ed) Comparative Genomics, vol 1&2. Humana Press, Totowa (NJ),

33. Jehl P, Sievers F, Higgins DG (2015) OD-seq: outlier detection in multiple sequence alignments. BMC bioinformatics 16 (1):1–11

34. Bailey TL, Johnson J, Grant CE, Noble WS (2015) The MEME Suite. Nucleic Acids Res 43 (W1):W39–W49. doi:10.1093/nar/gkv416

35. Anaconda software distribution. computer software. vers. 2-2.4. 0 (2015). Analytics Continuum,

36. Bezanson J, Karpinski S, Shah VB, Edelman A (2012) Julia: A fast dynamic language for technical computing. arXiv preprint 12095145

37. Jumper J, Evans R, Pritzel A, Green T, Figurnov M, Ronneberger O, Tunyasuvunakool K, Bates R, Žídek A, Potapenko A (2021) Highly accurate protein structure prediction with AlphaFold. Nature 596 (7873):583–589

38. Varadi M, Anyango S, Deshpande M, Nair S, Natassia C, Yordanova G, Yuan D, Stroe O, Wood G, Laydon A (2021) AlphaFold Protein Structure Database: massively expanding the structural coverage of protein-sequence space with high-accuracy models. Nucleic Acids Res

39. Sette M, van Tilborg P, Spurio R, Kaptein R, Paci M, Gualerzi CO, Boelens R (1997) The structure of the translational initiation factor IF1 from E. coli contains an oligomer-binding motif. The EMBO journal 16 (6):1436–1443

40. Reis AC, Salis HM (2020) An Automated Model Test System for Systematic Development and Improvement of Gene Expression Models. ACS Synthetic Biology 9 (11):3145–3156. doi:10.1021/acssynbio.0c00394

41. Keseler IM, Mackie A, Santos-Zavaleta A, Billington R, Bonavides-Martínez C, Caspi R, Fulcher C, Gama-Castro S, Kothari A, Krummenacker M (2017) The EcoCyc database: reflecting new knowledge about Escherichia coli K-12. Nucleic Acids Res 45 (D1):D543–D550

